# Microglia stabilize sleep homeostasis via adenosine A_3_ receptor signaling

**DOI:** 10.64898/2026.01.02.697450

**Authors:** Zhong Zhao, Xiaochun Gu, Jing Yu, Xueli Chen, Lifeng Zhang, Ting Zhao, Wei Ye, Heping Cheng

## Abstract

Sleep homeostasis maintains the sleep-wake balance through sleep pressure, a process orchestrated by the accumulation of extracellular adenosine (eADO). Microglial Ca²⁺ activity has been implicated in sleep regulation, but the mechanism whereby microglia sense sleep pressure remains unclear. Here we show that microglia regulate sleep homeostasis through brain state-dependent Ca^2+^ activity driven by adenosine A_3_ receptor (A_3_R) signaling. Using miniaturized two-photon microscopy (mTPM) in freely behaving mice, we demonstrate that microglial Ca²⁺ activity is rapidly altered by brain-state transitions. Pharmacological experiments reveal that microglial Ca^2+^ dynamics are predominantly mediated by A_3_R in response to brain state-dependent eADO oscillations. Microglia-specific deletion of A_3_R attenuates these state-dependent Ca^2+^ dynamics, impairs microglial morphological plasticity across sleep-wake cycles, and leads to sleep fragmentation by increasing transitions between wakefulness and non-rapid eye movement (NREM) sleep. Together, these findings establish that microglia regulate sleep homeostasis by stabilizing both wakefulness and NREM sleep, a process critically dependent on eADO-A_3_R signaling.

## Introduction

Sleep homeostasis which is driven by sleep pressure orchestrates the balance between sleep and wakefulness^1–3^. At the molecular level, it involves endogenous sleep-promoting substances (such as adenosine, TNF-α, and IL-1β), whose concentrations dynamically fluctuate with sleep pressure and contribute to the regulation of sleep^4–6^. Among these, extracellular adenosine (eADO) is a key molecule that progressively accumulates during wakefulness and drives sleep initiation^4,5^.

eADO exerts its physiological effects through four G-protein-coupled adenosine receptor subtypes: A_1_R, A_2A_R, A_2B_R, and A_3_R^7^. Among them, A_1_R and A_2A_R have been shown to modulate sleep in a brain region-specific manner^8,9^, whereas A_2B_R, a low-affinity receptor^10^, primarily responds to elevated eADO under pathological conditions^11^. In contrast, the physiological role of A_3_R in sleep regulation has not yet been clarified, despite its predominant expression in microglia.

Microglia, the resident immune cells of the central nervous system (CNS), are pivotal in regulating neural activity, synaptic pruning, and neuroimmunity^12,13^. Accumulating studies have established microglia as key modulators of sleep-wake states^14–18^, yet the cellular mechanisms underlying their responses to sleep pressure and subsequent homeostatic regulation remain to be elucidated. In their surveillance state, microglia continuously extend and retract their highly dynamic processes to monitor neural activity and maintain CNS homeostasis ^14,19–21^. Their dynamics are regulated by the purinergic signaling^22^. Through this pathway, P_2_Y_12_ receptors respond to ATP/ADP released from injured sites or neurons by inducing microglial process extension and morphological changes^23–25^. In this process, adenosine-activated A_3_R promotes ADP-induced extension of microglial processes^26^. Yet whether microglial A_3_R participates in physiological sleep regulation, and how adenosine signaling influences microglial dynamics across natural sleep-wake cycles, remain unknown.

Miniaturized two-photon microscopy (mTPM) has provided a transformative tool for *in vivo* imaging in freely behaving mice^27,28^, as evidenced by brain state-dependent microglial surveillance^14^. Specifically, it has been showed that microglial surveillance undergoes rapid and robust remodeling across natural sleep-wake cycles and in response to sleep deprivation stress through high-resolution time-lapse *in vivo* imaging in freely behaving mice^14^. Further, norepinephrine signaling modulates microglial surveillance, thereby serving as a key mechanism coupling microglia to brain state^14^. In the present study, we apply this powerful *in vivo* imaging approach to investigate the microglial signaling involved in regulating sleep homeostasis.

We aim to conduct long-term *in vivo* imaging of cortical microglial Ca^2+^ activity, morphology, and eADO levels in freely behaving mice using mTPM. By classifying brain states based on simultaneously recorded electroencephalogram (EEG) and electromyogram (EMG) signals, we intend to examine the relationship between sleep-wake states, eADO oscillations and microglial Ca^2+^ activity. Particular attention will be paid to elucidate the role of microglial A_3_R signaling as eADO-responsive elements. Together, these approaches will specific the role and mechanism of microglia as active modulators of sleep homeostasis.

## Results

### Microglial Ca^2+^ activity during natural sleep-wake cycles

To investigate how cortical microglial Ca²⁺ activity changes during natural sleep-wake cycles, we adopted a recently developed mTPM system (FHIRM-TPM) from our laboratory for long-term *in vivo* imaging in freely behaving mice^27,28^. Applying this system to a transgenic mouse model for microglia-specific GCaMP6s expression (*Cx3cr1^CreERT^; RCL-GCaMP6s*), we performed continuous *in vivo* imaging in the somatosensory cortex (Fig. 1a). By positioning the focal plane 50–100 μm beneath the cortical surface, we were able to stably record microglial Ca²⁺ dynamics at a high frame rate (5 Hz) over periods exceeding 6 hours. Throughout these prolonged recordings, the animals exhibited natural behavior and undisturbed sleep-wake cycles. To correlate the imaging data with brain states, EEG/EMG signals and behavioral video were simultaneously recorded for precise classification (Fig. 1a).

**Figure 1.**
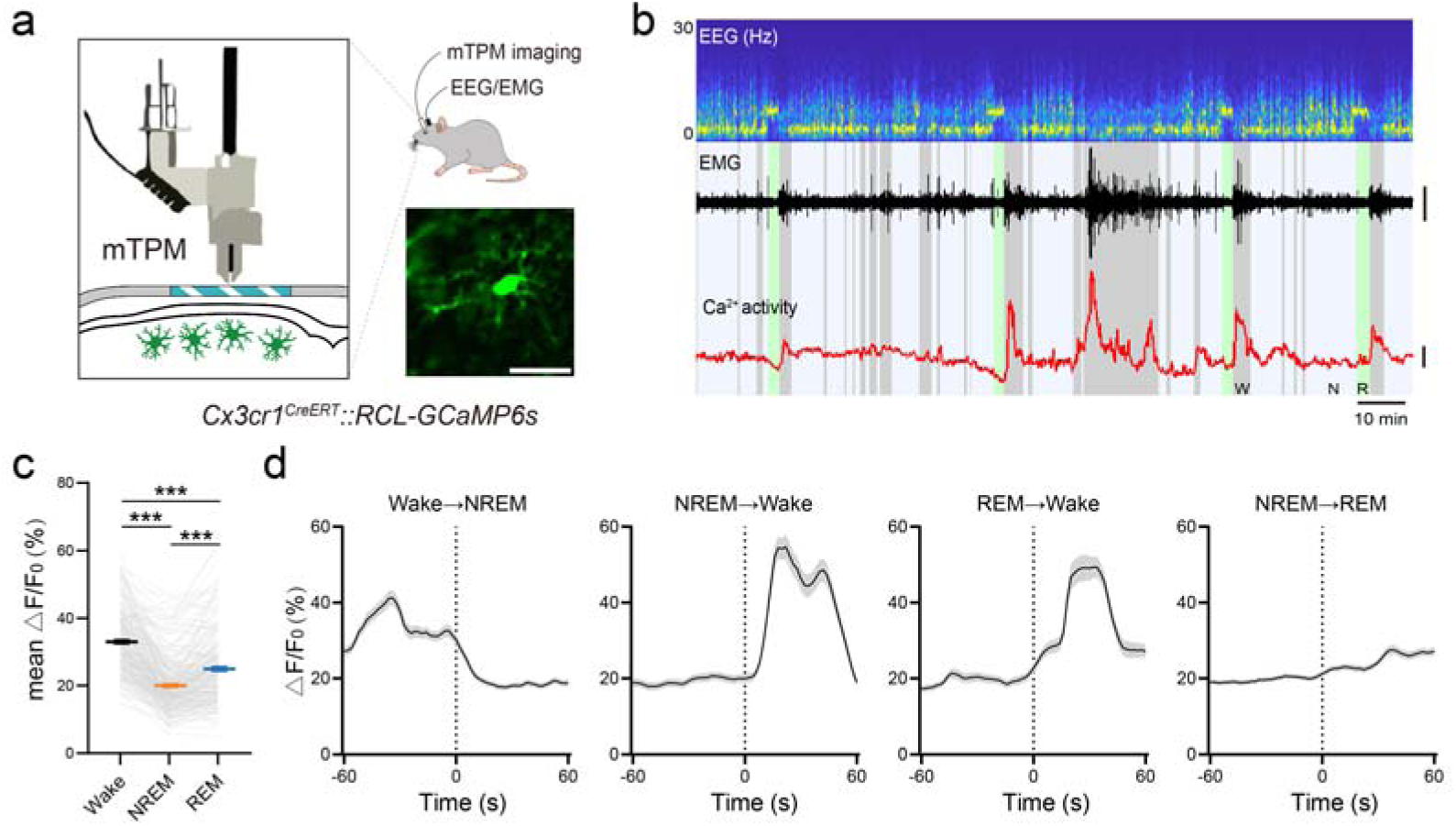
Brain state-dependent microglial Ca^2+^ activity. **a.** Experimental setup: mTPM and EEG/EMG recordings in freely behaving mice with tamoxifen-induced expression of GCaMP6s in microglia. A representative mTPM image of microglia in a *Cx3cr1^CreERT^::RCL-GcaMP6s* mouse. Scale bar, 30 μm. **b.** Representative long-term microglial Ca^2+^ activity during sleep-wake cycles in freely behaving mice, showing EEG (0–30 Hz spectrogram), EMG (scale bar, 100 μV), and z-scored Ca^2+^ signals from a microglia cell (scale bar, 2 z-score). Brain states are color-coded: wake (gray), NREM (purple), REM (green). **c.** Mean Ca^2+^ activity of microglia during different brain states. Friedman test with Dunn’s post-hoc test was used for paired measures. The gray solid line represents the calcium activity of the same cell across the three brain states. n = 220 cells from 19 sessions of 6 mice. \*\*\**p* < 0.001. **d.** Microglial Ca^2+^ activity during brain state transitions. The vertical dashed line represents the transition time. n = 143, 79, 61, and 164 cells from 10 sessions of 5 mice. Data are shown as mean ± s.e.m.

Microglial Ca²⁺ activity in freely behaving mice is enhanced during wakefulness (Fig. 1b). Furthermore, this activity was finely tuned by behavior, peaking during locomotion and dropping to significantly lower levels during quiet wakefulness and grooming, decreasing further during sleep (Fig. S1b). Beyond its behavioral patterns, the frequency of spontaneous Ca²⁺ transients was also modulated by sleep pressure, increasing with prolonged sleep deprivation and gradually returning to baseline during subsequent recovery sleep (Fig. S1c). Analysis across sleep-wake states revealed that microglial Ca²⁺ activity was highest during wakefulness, intermediate in rapid eye movement (REM) sleep, and lowest in NREM sleep (Fig. 1c). Notably, microglial Ca²⁺ activity shifted abruptly at transitions between wakefulness and NREM sleep, as well as from REM sleep to wakefulness, whereas the increase from NREM to REM sleep was only modest (Fig. 1d). Together, these results suggest that microglial Ca^2+^ activity exhibits robust, state-dependent dynamics during sleep-wake cycles, providing a foundation to investigate its functional role in the regulation of sleep homeostasis.

### Cortical eADO levels during natural sleep-wake cycles

The accumulation of eADO is a crucial factor driving sleep. In the cortex, eADO increases during sleep deprivation and gradually decreases during recovery sleep^29^, linking its dynamics to sleep homeostasis. To investigate how cortical eADO fluctuates across brain states, we expressed the adenosine sensor AAV9-hSyn-GRAB_ADO1.0_ in the somatosensory cortex and performed long-term *in vivo* imaging in freely behaving mice using mTPM, while simultaneously monitoring brain states via EEG/EMG recordings (Fig. 2a).

**Figure 2.**
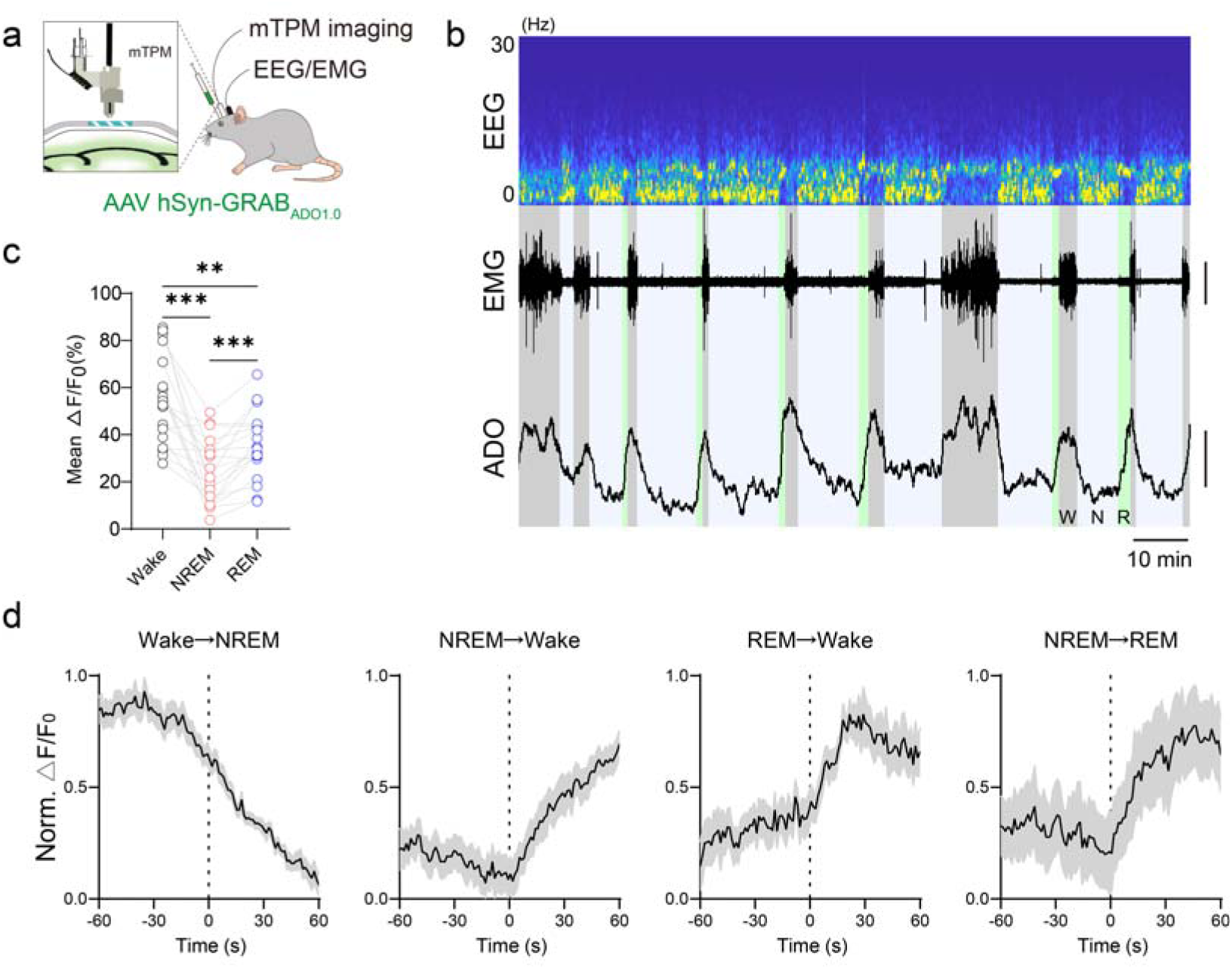
eADO oscillations during natural sleep-wake cycles. **a.** Schematic diagram depicting mTPM images of eADO reported by the GRAB_ADO1.0_ sensor expressed in neurons. Freely behaving mice were head-mounted with a mTPM and EEG/EMG electrodes. **b.** Representative simultaneous recordings from somatosensory cortex during the sleep-wake cycles in freely behaving mice, showing EEG (0–30 Hz spectrogram), EMG (scale bar, 1 mV), and eADO (scale bar, 2 z-score) signals. **c.** Mean eADO levels in different brain states. Data from the same recording are connected by lines. One-way ANOVA with Tukey’s post-hoc test was used. n = 18 sessions from 5 mice. \*\**p* < 0.01, \*\*\**p* < 0.001. **d.** Signal of the GRAB_ADO_ sensor during brain state transitions. The vertical dashed line represents the transition time. n = 30, 30, 13, and 17 events from 4 mice. ns, not significant. Data are shown as mean ± s.e.m.

We found that eADO displayed sustained high levels during wakefulness, declined to its lowest levels in NREM sleep, and was marked by brief, rapid increases during REM sleep (Fig. 2b). Quantification confirmed that eADO levels were highest during wakefulness, intermediate during REM sleep, and lowest during NREM sleep (Fig. 2c). Furthermore, eADO responded rapidly to brain state transitions, decreasing during the transition from wakefulness to NREM sleep, but increasing sharply during transitions from NREM sleep to either wakefulness or REM sleep, and again from REM sleep to wakefulness (Fig. 2d). Together, these results indicate that cortical eADO dynamics are closely coupled to sleep-wake transitions.

### eADO-A_3_R regulates microglial Ca^2+^ activity

Adenosine regulates Ca^2+^ activity through multiple adenosine receptors, with A_3_R being the predominantly expressed adenosine receptor in microglia^22,26^. To investigate whether microglial Ca^2+^ activity is mediated by the eADO-A_3_R axis, we performed intracerebral administration of adenosine, an A_3_R agonist (Cl-IB-MECA), or an A_3_R antagonist (MRS1523) and assessed their effects on microglial Ca^2+^ activity (Fig. 3a). Given that drug administration might alter microglial motility, Ca²⁺ events were quantified using Astrocyte Quantitative Analysis (AQuA) software to minimize movement-related artifacts. Intracerebral administration of ACSF did not change Ca^2+^ activity (Fig. 3b; Fig. S2a), allowing us to track the dynamic pharmacological effects within the same field of view (FOV). Both adenosine and Cl-IB-MECA increased the number of Ca^2+^ events (Fig. 3c-e), whereas MRS1523 decreased event frequency (Fig. 3f, g). Neither Cl-IB-MECA nor MRS1523 altered the amplitude of microglial Ca²⁺ activity (Fig. 3e, g).

**Figure 3.**
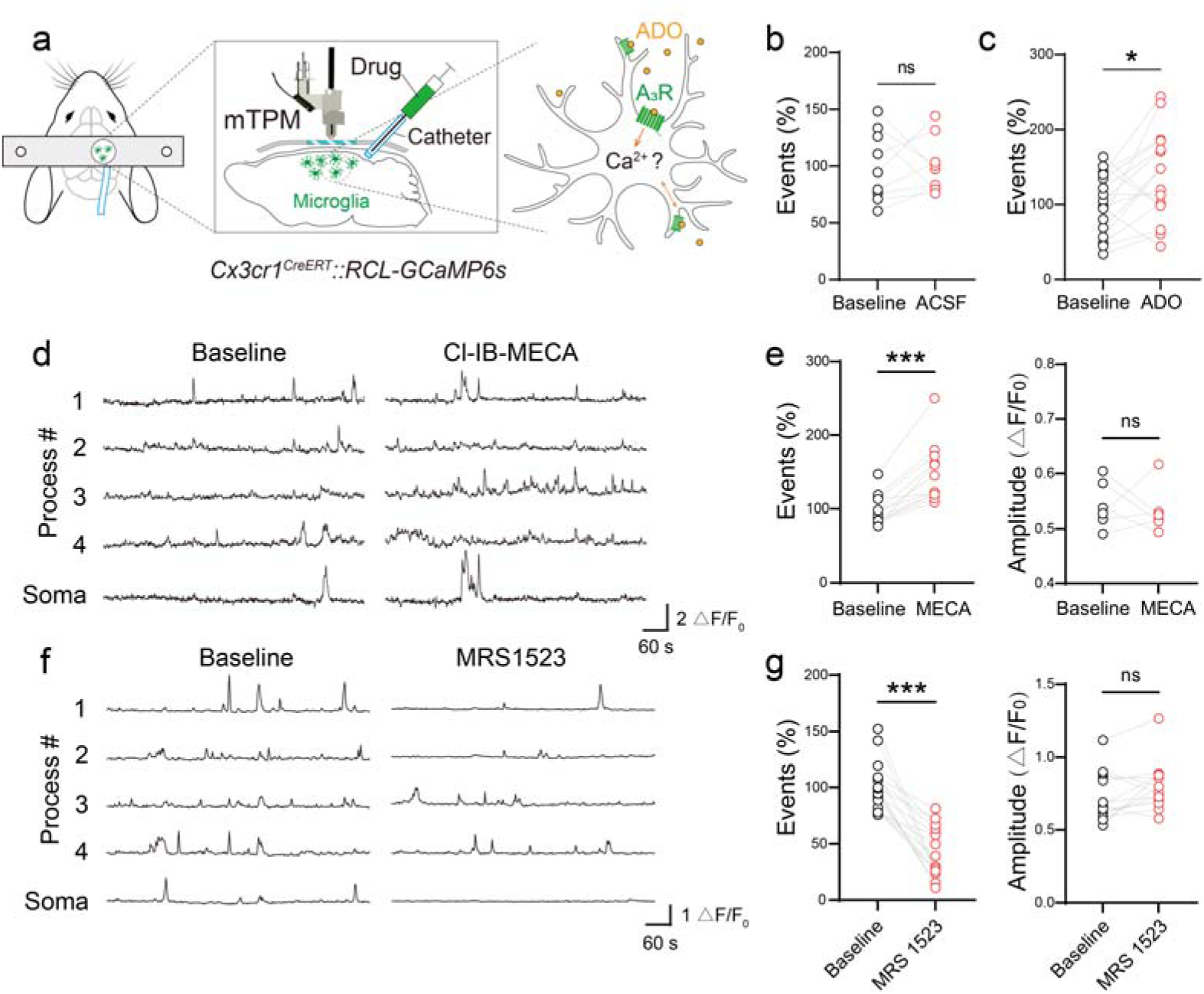
Microglial Ca²⁺ activity is modulated by ADO and A_3_R. **a.** Schematic diagram depicting mTPM images of microglial Ca^2+^ activity and morphology in freely behaving mice. A drug infusion catheter was implanted in the subarachnoid space adjacent to the cranial window. Microglia were imaged following local application of ADO or A_3_R ant/agonists for 30 min. **b–c.** The number of Ca^2+^ events within the FOV in 5-minute recordings, 30 min after ACSF (**b**) or ADO (**c**) administration. Data from the same FOV are connected by lines. *n* = 9 FOVs from 3 mice (ACSF). *n* = 16 FOVs from 5 mice (ADO). **d**, **f.** Representative microglial Ca^2+^ signal in 10-min recordings, 30 min after Cl-IB-MECA (**d**) or MRS1523 (**f**) administration. **e**, **g.** The number of Ca^2+^ events and amplitude in microglia processes in 5-minute recordings, 30 min after Cl-IB-MECA (**e**) or MRS1523 (**g**) administration. Data from the same FOV are connected by lines. \**p* < 0.05, \*\*\**p* < 0.001. ns, not significant.

To test whether these effects were directly mediated by microglial A_3_R, we generated microglia-specific A_3_R knockout mice by crossing *adora3^fl/fl^* (*adora3* cKO) mice with *Cx3cr1^CreERT^* mice (Fig. S2b). Tamoxifen was administered to 7-week-old mice to ablate *adora3* in microglia. qRT-PCR analysis of isolated CD11b^+^/CD45^low^ microglia showed a ∼75% reduction in *adora3* mRNA in A_3_R cKO mice, with no significant change in the negative (CD11b^-^/CD45^-^) cell population (Fig. S2c, d). The regulatory effects of Cl-IB-MECA and MRS1523 were significantly attenuated in these mice (Fig. S2e, f), indicating that microglial A_3_R directly regulates Ca^2+^ activity. Collectively, these findings establish the eADO-A_3_R pathway as a crucial regulator of microglial Ca²⁺ activity, enabling microglia to respond dynamically to eADO fluctuations across brain states.

### Microglial A_3_R deficiency fragmentizes wakefulness and NREM sleep

To explore the role of microglial A_3_R in sleep regulation, we monitored the sleep-wake patterns in microglial A_3_R cKO mice. Sleep-wake states analysis using EEG/EMG recordings (ZT3–ZT9 and ZT15–ZT21) revealed no changes in the total duration of wakefulness, NREM, or REM sleep (Fig. 4a, b). However, microglial A_3_R cKO mice exhibited more frequent brain state transitions (Fig. 4a, c). Further analysis revealed more frequent transitions between wakefulness and NREM sleep (Fig. 4a, d), as both states were fragmented into shorter episodes (Fig. 4e, f).

**Figure 4.**
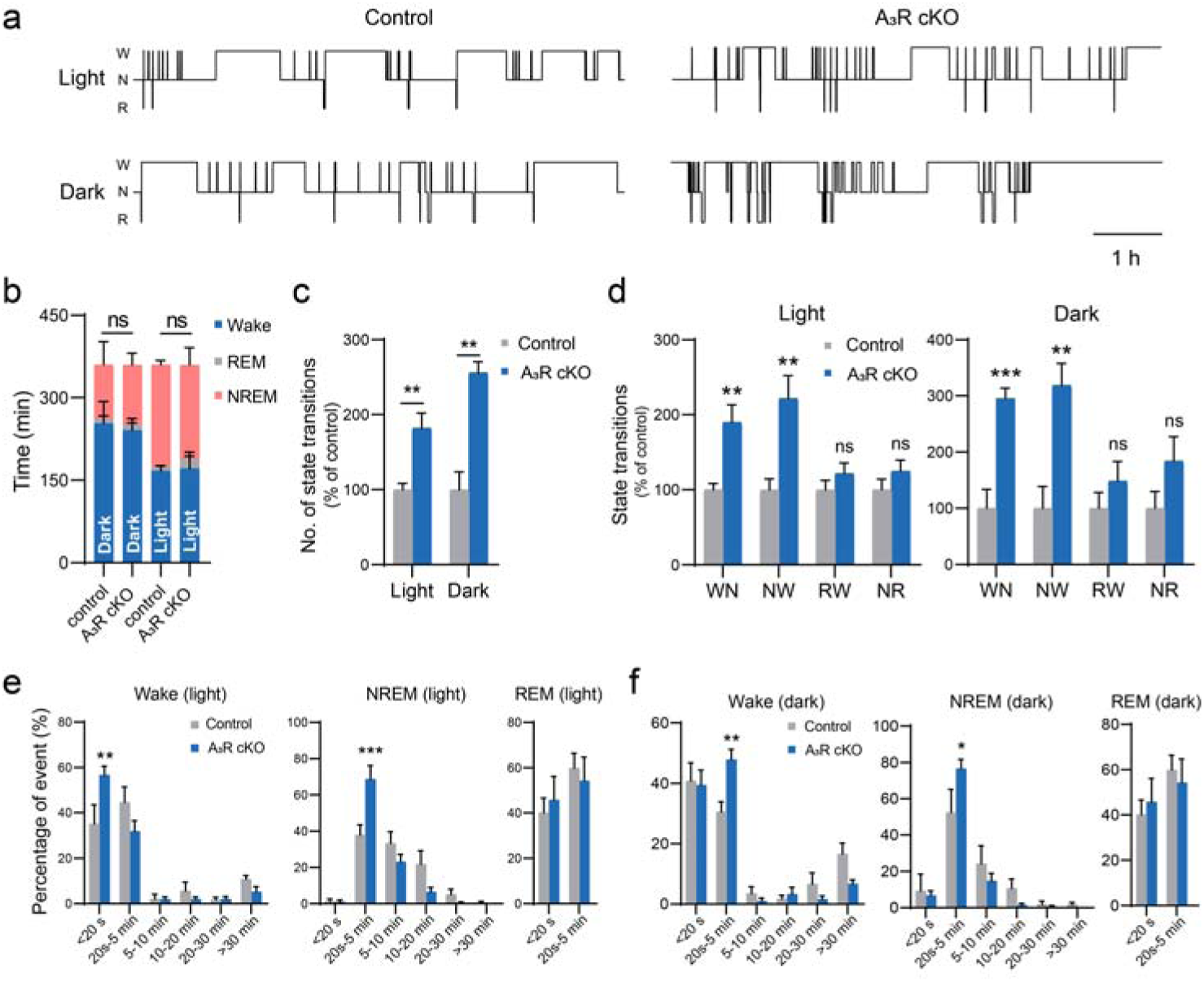
Sleep fragmentation in microglial A_3_R cKO mcie. **a.** A hypnogram measured from microglia A_3_R cKO and sibling control mice at the light phase (ZT3-9) and dark phase (ZT15-21). *Cx3cr1^CreERT^::Adora3^fl/fl^* mice, microglial A_3_R cKO mice; *Cx3cr1^WT^::Adora3^fl/fl^* mice served as littermate controls. **b.** Total duration of wakefulness, NREM sleep, and REM sleep at the light phase (ZT3–9) and dark phase (ZT15–21) in control and A_3_R cKO mice. n = 5 mice per group. Two-way ANOVA with post-hoc comparison between control and A_3_R cKO mice. **c.** Changes in the number of brain state transitions in control and A_3_R cKO mice. n = 5 mice per group. Unpaired t test was used. **d.** Number of four brain state transitions in control and A_3_R cKO mice. Unpaired t test was used. n = 5 mice per group. **e-f.** Percentage of episode duration of wakefulness, NREM sleep, and REM sleep in control and A_3_R cKO mice. Two-way ANOVA with post-hoc comparisons test was used. n = 5 mice per group. \*\**p* < 0.01, \*\*\**p* < 0.001. ns, not significant. Data are shown as mean ± s.e.m.

Microglial A_3_R cKO mice exhibited a higher proportion of brief microarousals (< 20 s) in the light phase (Fig. 4e) and short arousals (20 s–5 min) in the dark phase (Fig. 4f). The proportion of short NREM sleep also increased during light/dark phases, while REM architecture remained stable (Fig. 4e, f). These results indicate that microglial A_3_R signaling contributes to maintaining the stability of wakefulness and NREM sleep.

### Microglial A_3_R regulates state-dependent Ca^2+^ activity during natural sleep-wake cycles

To further test the eADO-A_3_R-Ca^2+^ axis of sleep regulation, we next investigated whether microglial A_3_R modulates state-dependent Ca^2+^dynamics during natural sleep-wake cycles in freely behaving mice. To this end, we generated *Cx3cr1^CreERT^::adora3^fl/fl^::RCL-GCaMP6s*mice, in which tamoxifen induces microglial-specific knockout of *adora3* and simultaneous expression of GCaMP6s.

Compared with *Cx3cr1^CreERT^::RCL-GCaMP6s* control mice, microglial Ca^2+^ activity in A_3_R cKO mice was significantly decreased during wakefulness and REM sleep, but increased during NREM sleep (Fig. 5a). Consequently, the differences in microglial Ca²⁺ activity across wakefulness and sleep were attenuated (Fig. 5a, b). Further analysis showed that, similar to controls, A_3_R cKO mice exhibited higher microglial Ca²⁺ activity during wakefulness than during sleep (Fig. 2c, 5a). However, microglial Ca^2+^ activity did not differ significantly between NREM and REM sleep in A_3_R cKO mice (Fig. 5b).

**Figure 5.**
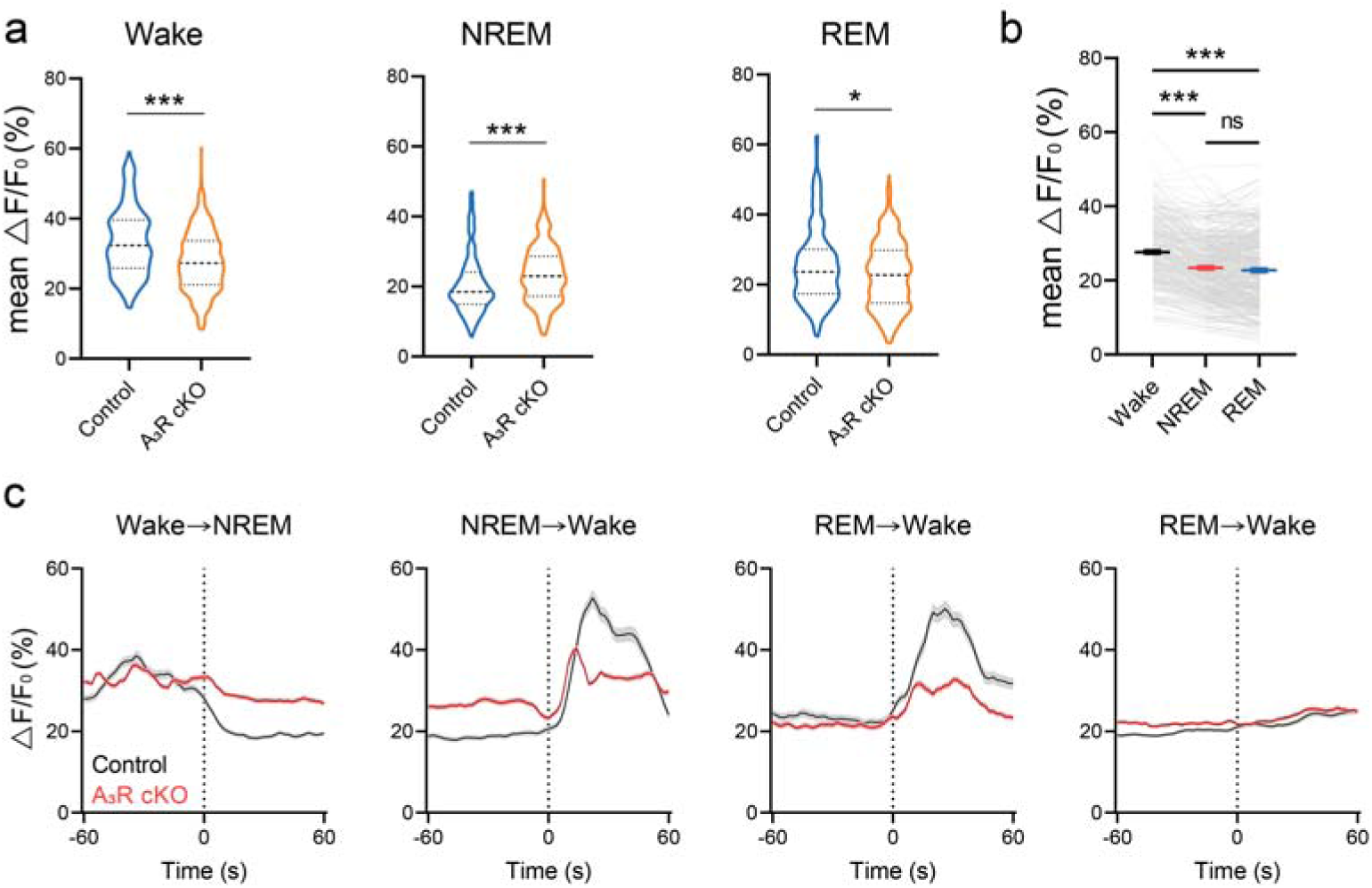
A_3_R regulates state-dependent microglial Ca^2+^ activity. **a.** Mean Ca^2+^ activity of microglia during different brain states in littermate control and A_3_R cKO mice. Control mice, *Cx3cr1^CreERT^::RCL-GCaMP6s* mice; A_3_R cKO mice, *Cx3cr1^CreERT^::Adora3^fl/fl^::RCL-GCaMP6s* mice. Experimental setup: *In vivo* imaging was performed one month after tamoxifen induction. Mann-Whitney test was used. n = 220 cells from 19 sessions of 6 control mice, n = 372 cells from 15 sessions of 5 A_3_R cKO mice. **b.** Mean Ca^2+^ activity of microglia in A_3_R cKO mice during different brain states. Data from the same cell are connected by lines. Friedman test with Dunn’s post-hoc test was used. n = 372 cells from 15 sessions of 5 mice. **c.** Ca^2+^ activity during brain state transitions in control and microglial A_3_R cKO mice. The vertical dashed line represents the transition time. n = 220 cells from 19 sessions of 6 control mice, n = 372 cells from 15 sessions of 5 A_3_R cKO mice. n = 144, 80, 62, and 164 cells from 5 control mice. n = 620, 540, 360, and 686 cells from 5 A_3_R cKO mice. \**p* < 0.05, \*\*\**p* < 0.001. ns, not significant. Data are shown as mean ± s.e.m.

Although microglial Ca²⁺ activity in A_3_R cKO mice still followed the general trend of responding to brain state transitions, the amplitude of these transitions was significantly reduced (Fig. 5c). Collectively, these findings indicate that microglial A_3_R is essential for maintaining state-dependent Ca²⁺ activity and for amplifying Ca²⁺ dynamics during transitions, thereby enabling microglia to synchronize their activity with ongoing sleep-wake cycles. The attenuation of wake-sleep Ca^2+^ change and the loss of NREM-REM Ca^2+^ difference may underlie the more frequent brain state transitions between wakefulness and NREM sleep.

### A_3_R regulates microglial morphology during natural sleep-wake cycles

Our previous work demonstrated that microglial morphology changes are brain state-dependent and modulated by sleep need^14^. In cultured primary microglia, A_3_R promoted ADP-induced microglial process extension^26^, suggesting a role for A_3_R in mediating microglial morphological plasticity. Since microglial morphology is closely linked to intracellular Ca²⁺ signaling, we next asked whether A_3_R also regulates microglial morphology across natural sleep-wake cycles.

To address this, we generated *Cx3cr1^CreERT^::adora3^fl/fl^::Cx3cr1^GFP/-^* mice, in which tamoxifen induces microglial-specific knockout of *adora3* in GFP-labeled microglia (Fig. 6a). In control mice without gene knockout, microglial processes exhibited dynamic changes in response to brain state (Fig. 6a-d). Specifically, microglial process length, territory area, and branch point number were greater during NREM sleep than during wakefulness or REM sleep, and lowest during wakefulness (Fig. 6b-d). This result is consistent with our previous findings, confirming that microglial morphology is dynamically remodeled by sleep state. However, in microglial A_3_R cKO mice, these morphological differences across brain states were abolished. The length, area, and branch point number of microglial processes during wakefulness were increased compared to those in control mice (Fig. 6a-d), demonstrating a disruption of normal state-dependent regulation. Together, these results indicate that microglial A_3_R is essential for state-dependent morphological and functional changes in microglia across natural sleep-wake cycles, highlighting the role of adenosine signaling in microglial dynamics.

**Figure 6.**
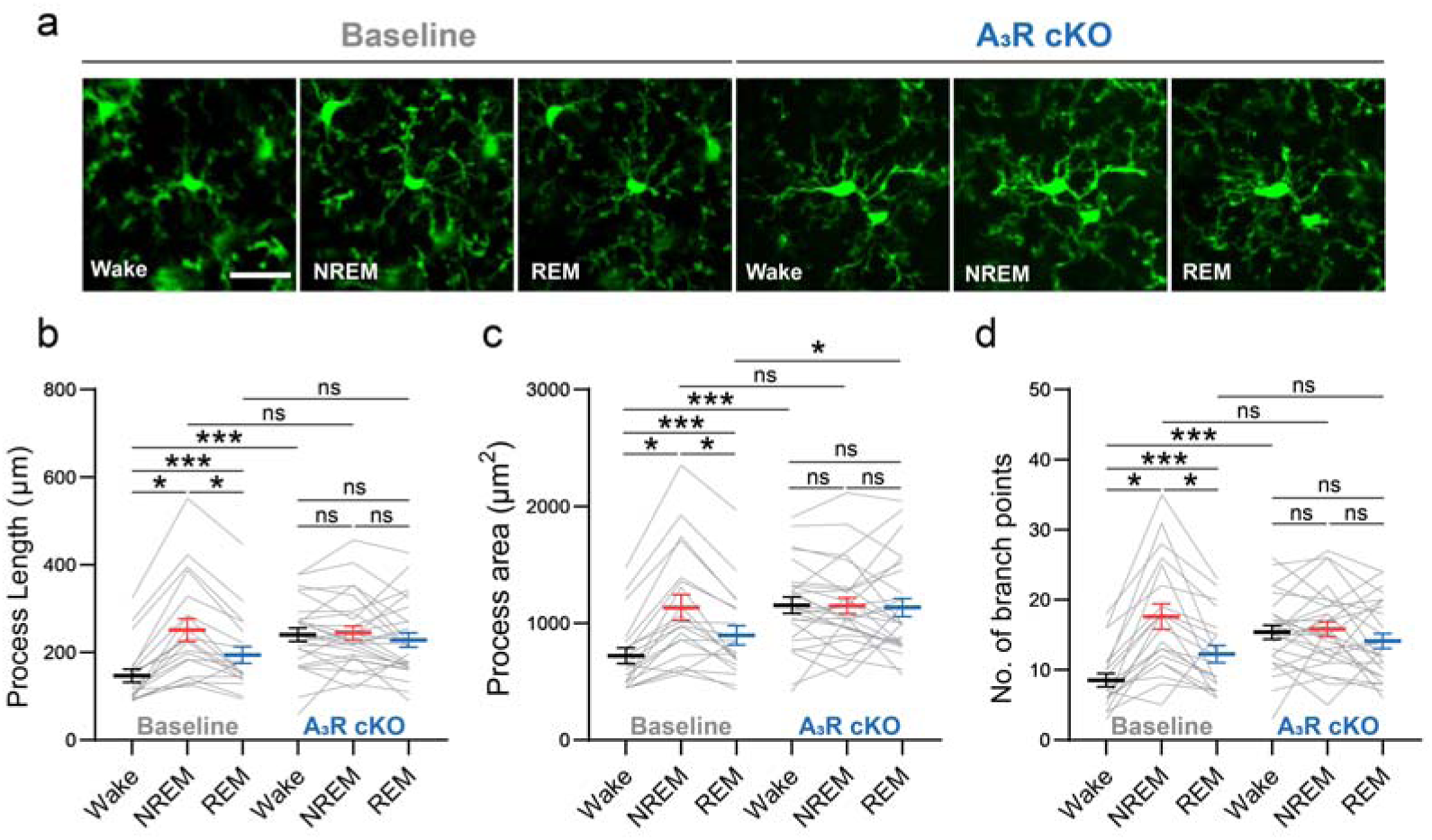
A_3_R alters state-dependent morphological dynamics in microglia. **a.** Representative microglial morphological changes during the sleep-wake cycle in microglial A_3_R cKO mice (*Cx3cr1^CreERT^::Adora3^fl/fl^::Cx3cr1^GFP/-^*mice). Data collection was performed as follows: baseline data were acquired at 4 weeks post-surgery, tamoxifen was induced at 5 weeks, and A_3_R cKO data were collected at 9 weeks post-surgery. Scale bar, 30 μm. **b–d.** Quantification of microglia processes length (**b**), area (**c**), and number of branch points (**d**) before (baseline) and after (A_3_R cKO) tamoxifen-induced microglial A_3_R conditional knockout. Data from the same microglia are connected by lines. Friedman test with Dunn’s post-hoc test (baseline: length and area) and one-way ANOVA with Tukey’s post-hoc test (baseline: branch points; A_3_R cKO: length, area and branch points) were used. Data are from 5 mice (Baseline, n = 20 cells; A_3_R cKO, n = 27 cells). \**p* < 0.05, \*\*\**p* < 0.001. ns, not significant. Data are shown as mean ± s.e.m.

## Discussion

In this study, leveraging mTPM in freely behaving mice, we delineate the role of microglial eADO-A_3_R-Ca^2+^ signaling axis in sleep homeostasis. We demonstrate that cortical eADO levels and microglial Ca²⁺ activity exhibit brain state-dependent dynamics, co-varying with sleep pressure. Critically, we identify microglial A_3_R as the key receptor transducing fluctuating eADO levels into intracellular Ca²⁺ signals, and establish that disruption of this axis fragmentizes sleep-wake architecture and abolishes state-dependent microglial morphological plasticity (Fig. 7). These findings collectively define a novel purinergic-immune pathway wherein microglia, via A_3_R, act as dynamic sensors of sleep pressure to regulate sleep stability.

**Figure 7.**
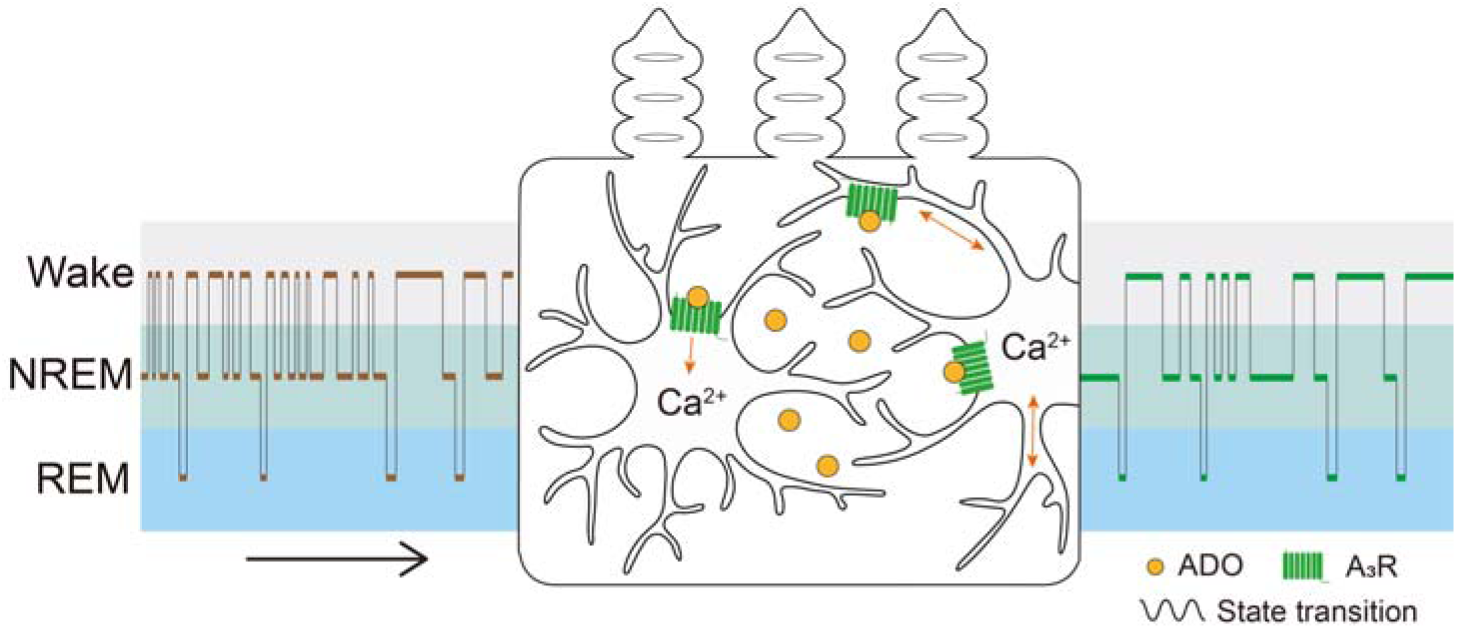
Microglia as stabilizers of sleep homeostasis. Regulation of A_3_R-mediated microglial Ca^2+^ activity and morphology dynamics by eADO oscillations underlies the stabilization of wake-sleep states in mice. Microglia serve as a stabilizer for sleep homeostasis through the eADO-A_3_R signaling pathway.

Adenosine receptors predominantly regulate sleep/wake states by modulating neuronal activity and neurotransmitter systems^9^. Beyond the well-characterized roles of neuronal adenosine receptors (A_1_R and A_2A_R) in sleep regulation^30–36^, we identify microglial A_3_R as a pivotal non-neuronal mediator of sleep homeostasis. Previous studies showed that constitutive knockout of A_3_R increased nocturnal activity in female mice in the dark and reduced behavioral activation in drug (caffeine, amphetamines) stimuli^37^. We show that microglial A_3_R maintains state-dependent Ca²⁺ activity and morphological dynamics, thereby curbing transitions between wakefulness and NREM sleep and promoting the stability of both states. Notably, as a Gi/Gq_/11_-coupled protein receptor^7^, A_3_R induces intracellular Ca^2+^ accumulation and inhibits adenylyl cyclase activity, suggesting that microglial A_3_R may modulate Ca^2+^ activity through coordinated cAMP-dependent signaling to influence sleep-wake states.

The disruption of the eADO-A_3_R-Ca²⁺ signaling axis in our study leads to a defined sleep phenotype characterized by fragmentation of wakefulness and NREM sleep, without affecting total sleep duration or REM sleep dynamics (Fig. 4). This "instability" phenotype, marked by increased state transitions and shorter wake/NREM bout durations, coincides with attenuated state-dependent fluctuations in microglial Ca²⁺ activity and morphological plasticity in A_3_R cKO mice (Fig. 5, 6), highlighting the axis’s role in maintaining sleep-wake stability rather than gross sleep quantity. This phenotype aligns with a growing body of literature indicating that microglia are important for NREM sleep stability and architecture. Global microglial depletion models consistently report increased NREM sleep fragmentation^15–17^, often accompanied by alterations in total NREM sleep time^15,17^—a more pronounced effect than the specific signaling disruption observed in our A_3_R cKO model. This distinction underscores that while the presence of microglia broadly supports sleep homeostasis, specific signaling pathways, such as the eADO-A_3_R-Ca²⁺ signaling axis, fine-tune particular aspects of state stability. Our findings thus refine the understanding of microglial sleep regulation by revealing an adenosine-mediated pathway through which microglia govern the consolidation of wakefulness and NREM sleep, likely by modulating neural circuit excitability at state transition points.

Beyond the potential impact of distinct waking and sleep states on microglial dynamics (Fig. 1), prior studies have elucidated that the attributes of these dynamics vary across both subcellular compartments and brain regions^18,38,39^. Throughout the sleep-wake cycle, microglial morphology also exhibits region-specific traits^39^. For example, *in vitro* brain slice recordings revealed that somatosensory cortical microglia occupy smaller territories than those in the hippocampus or basal forebrain, with morphological changes related to slow-wave activity intensity^39^. Consistent with this, our prior findings demonstrated that microglial morphology in the somatosensory cortex is brain-state-dependent^14^. Functionally, chemogenetic activation of Gi protein-mediated microglial Ca^2+^ activity in the prefrontal cortex or basal forebrain modifies the sleep/wake states, while similar manipulations in the dorsal striatum do not^18^. This indicates that the influence of microglial Ca^2+^ activity on sleep regulation varies across different brain regions. Moreover, we observed that microglial Ca^2+^ activity peaked during wakefulness in the somatosensory cortex of freely behaving mice (Fig. 1), whereas it was reported to be higher during NREM sleep in the prefrontal cortex of head-fixed mice^18^. This suggests that microglial Ca^2+^ activity is region-specific across the brain during the sleep-wake cycle, and the variations in local neuronal activity may be a key driver of these distinct patterns.

In summary, our study establishes microglial A_3_R as a central mediator of Ca^2+^ dynamics and morphology during natural sleep-wake cycles. By decoding brain-state-specific fluctuations in eADO, microglial A_3_R consolidates sleep architecture by reducing state-transition frequency and preserving stability, thereby enforcing sleep homeostasis. Future studies should elucidate the molecular pathways downstream of A_3_R and investigate how microglia interact with other sleep-regulating cells, including neurons and astrocytes. As demonstrated here, 24-h continuous *in vivo* imaging and multimodal recording in freely behaving animals would empower neuroscientists to gain novel insights into these important questions.

## Methods

### Animals

All experimental procedures were conducted following the approval of the Institutional Animal Care and Use Committee of PKU-Nanjing Institute of Translational Medicine (Approval ID: IACUC-2021-023). C57BL/6 mice aged 2–3 months were used in most of the experiments, with a few longer data collection periods using mice at around 4 months of age. Mice were housed with a standard 12h light/12h dark cycle and fed standard chow *ad libitum*.

*Cx3cr1^CreERT^* (#020940) mice and *Cx3cr1^GFP/GFP^*(#005582) mice were purchased from the Jackson Laboratory. *Adora3^fl/fl^*(#T006284, *adora3* cKO) mice were purchased from the GemPharmatech Co., Ltd. *RCL-GCaMP6s* mice were provided by Nanjing Raygenitm Biotech Co., Ltd. *Cx3cr1^CreERT^* mice and *adora3^fl/fl^* mice were crossed to generate the conditional knockout mouse line.

To generate mice with microglia-specific expression of GCaMP6s and A_3_R knock-out, we bred *RCL-GCaMP6s::adora3^fl/-^*mice with *Cx3cr1^CreERT^::adora3^fl/-^* mice to obtain *Cx3cr1^CreERT^:: RCL-GCaMP6s::adora3^fl/fl^* mice. Similarly, in order to generate mice with microglia-specific expression of GFP and A_3_R knock-out, we first bred *Cx3cr1^GFP/GFP^::adora3^fl/-^*mice with *Cx3cr1^CreERT^::adora3^fl/-^* mice to obtain *Cx3cr1^CreERT^::Cx3cr1^GFP/-^::adora3^fl/fl^*mice. To activate tamoxifen-induced CreERT, mice were intraperitoneally injected at ∼8 weeks of age with 100 μL 20 mg/mL tamoxifen (T5648, Sigma) for five consecutive days, and considering the recovery period of cranial window surgery and the macrophage turnover, mice that had been induced with tamoxifen one month prior were selected for data acquisition.

### Surgery

All surgical interventions were performed aseptically, with all substances administered to the animals being sterile. The mice were anesthetized using isoflurane (1.5% in air at a flow rate of 0.5 L/min) and kept on a 37°C heating pad throughout the surgery. The region of the cerebral cortex to be imaged was identified based on stereotactic coordinates (somatosensory area, from bregma: anteroposterior, -1 mm, mediolateral, +2 mm). A 4-mm cranial window was created using a high-speed microdrill under the guidance of an anatomical microscope. Glass coverslips were implanted and fixed onto the somatosensory cortex. Mice were used after 4 weeks of postoperative recovery.

Implantation of epidural screw electrodes (with a diameter of 0.8 mm) was performed at opposite sides of the imaging window to enable continuous EEG recording from the frontal cortex (bregma: anteroposterior, +1.5 mm; mediolateral, +1.5 mm) and parietal cortex (anteroposterior, -2 mm; mediolateral, +2.5 mm). The electrodes were fixed to the skull using dental cement. Additionally, another set of electrodes was inserted into the neck muscles for EMG recording. Subsequently, a headpiece baseplate was affixed to the skull using cyanoacrylate and reinforced with dental cement. Mice were used after at least two weeks of postoperative recovery.

### Virus injection

200 nL of AAV9-hsyn-GRAB_ADO1.0_ (PT-1348, BrainVTA) was injected into the somatosensory cortex (A/P, -1 mm; M/L, +2 mm; D/V, -0.3 mm) at a rate of 20 nL/min. A 4-mm-diameter glass coverslip was implanted and fixed on the somatosensory cortex. The titer of the NE and ADO sensor viruses was estimated to be ≥ 2×10^12^ vg/mL. The mice were used one month after postoperative recovery.

### Two-photon imaging and sleep recording

Mice recovered from surgery without cranial window infection and with typical EEG/EMG signals were selected for imaging. Initial imaging stacks were acquired to select the best FOV using mTPM (Transcend Vivoscope) on head-fixed mice^27,28^. For Ca^2+^ activity imaging, suitable FOVs with rich Ca^2+^ activity of microglia processes were selected at a depth of 50–100 μm below the dura mater. For microglial morphology imaging, appropriate FOVs containing 5–10 microglial somata were selected. To achieve multi-day imaging of the same FOV and repeated mounting, the mTPM holder was sealed onto the headpiece baseplate positioned above the coverslip.

FHIRM-TPM was used to collect imaging data^27,28^. Imaging of Ca^2+^ activity and morphology was performed in high-resolution model (lens NA 0.7, FOV 220 μm × 220 μm, lateral resolution 0.74, axial resolution 6.53 μm) combined with ETL model^27,28^. The sampling frame rate for Ca^2+^ activity and morphology imaging was 10–14 Hz and 5–7 Hz, respectively. In addition, large FOV model (lens NA 0.75 FOV 420 μm × 420 μm, lateral resolution 1.13, axial resolution 12.2 μm) was used in adenosine imaging. The sampling frame rate was 5 Hz.

The continuous and simultaneous recording of EEG/EMG signals was performed using the Vital Record system developed by Kissei Comtec, while mouse behavior was monitored with an infrared video camera. Mice utilized for imaging and recording purposes were housed in the behavioral area for three days prior to the experiment for acclimation. During this period, they were allowed unrestricted access to food and water.

### Drug application *in vivo*

A drug infusion catheter filled with ACSF was placed in the subarachnoid space adjacent to the imaging area of the cranial window. The catheter and the skull were held together using biological tissue glue and dental cement, and then the headpiece baseplate was affixed to the skull using cyanoacrylate and reinforced with dental cement. Mice were used after at least two weeks of postoperative recovery. During the administration process, the needle of the microsampler was positioned deep within the catheter and close to the opposite end of the catheter. A total volume of 10 μL of solution was delivered through the catheter and into the brain. Durgs were applied at the following concentrations: 500 nM adenosine (ST1075, Beyotime Biotechnology), 300 nM Cl-IB-MECA (C277, Sigma), 1 μM MRS1523 (M1809, Sigma) or ACSF. Ca^2+^ activity and morphology of microglia were recorded before and 30 minutes after administration.

### Sleep deprivation

Mice were deprived of sleep for 6 hours using a sleep deprivation device (XR-XS108, Shanghai Xinsoft Information Technology). The parameters were set as follows: running direction of the bar, positive and negative rotation; stripping strength, moderate; rotating speed, 5 rpm; running mode, random.

### Microglia isolation from adult mice

Cortical microglia were isolated from A_3_R cKO and control mice one month after tamoxifen induction. Anesthetized mice were perfused prechilled PBS to obtain brain samples. The cerebral cortex was dissected, thoroughly chopped, and then digested with 200 UI of papain (P3125, Sigma-Aldrich) at 37°C for 30 minutes, followed by termination of the digestion with a medium containing 10% fetal bovine serum (18045088, Gibco). The solution was filtered through a 70-μm cell strainer to remove debris. The cell suspension was centrifuged at 300 g for 5 minutes, and the cell pellets were resuspended in 4 mL of 30% Percoll plus (P331004, Aladdin), transferred to a 15 mL centrifuge tube containing 4 mL of 70% Percoll plus, on which 2 mL of PBS was layered, and then centrifuged at 18°C for 40 minutes, and the top layer of myelin was removed. The cell pellets were washed twice with fluorescent activated cell sorting (FACS) buffer, and CD11b (101211, Biolegend) and CD45 (157213, Biolegend) immunostaining was performed on ice. Microglia and negative cells were sorted in a flow cytometer (FACSAria II, Becton Dickinson). After FACS, A_3_R expression level was detected by mRNA isolation and real-time quantitative PCR.

### Quantitative real-time PCR

Total RNA was extracted from primary cells enriched by centrifugation, according to the guidelines of RaPure cell RNA kit (R4010, Magen). First-strand cDNA was synthesized using M-MLV Reverse Transcriptase kit (R223, Vazyme). Quantitative real-time PCR was conducted using a Hieff qPCR SYBR Green Master Mix (No Rox) kit (11201ES03, Yeasen). The mRNA level of β*-actin* was analyzed as an internal control. The following primer sequences were used: *adora3* forward primer: ACATGCTGGTCATCTGGGTG; *adora3* reverse primer: AGCAGCACACAGGACATGAA; β*-actin* forward primer: TTGGGTATGGAATCCTGTGGC; β*-actin* reverse primer: GCTAGGAGCCAGAGCAGTAATC.

### Immunofluorescence

Brain slices fixed with 4% paraformaldehyde were permeabilized by treating with 0.5% TritonX-100. Following a 2-hour blocking period in PBS containing 2% BSA at 37°C, brain slices were incubated overnight at 4°C with rabbit-derived primary antibody against IBA1 (17198, Cell Signaling Tech), GFAP (80788, Cell Signaling Tech), or NeuN (24307S, Cell Signaling Tech). Slices were then incubated in the dark with a fluorescently labeled secondary antibody for 2 hours at 37°C and washed with PBS. Finally, slices were sealed with an anti-fluorescent quench agent containing DAPI.

### Quantitative analysis

#### Microglial Ca^2+^ activity analysis

The time-lapse stack was aligned offline using the Template-Matching plugin of ImageJ Fiji software to correct for xy motion artifacts. To analyze the changes of long-term Ca^2+^ activity in microglia processes in freely behaving mice, ImageJ Fiji was used to delineate the region of interest (ROI) and extract the data, which were then processed in MATLAB (MathWorks). Raw data were binned into 1Hz, followed by background autofluorescence subtraction. ΔF/F_0_ was subsequently calculated, utilizing the fluorescence signal itself. F_0_ was defined as the average value of the lower 25th quartile fluorescence signal within each 60-second window of the recorded fluorescence signal^38^.

In the experiment of drug application, since drug stimulation accelerates the movement of microglial processes, we used AQuA software^40^, using spatially unfixed and size-varying of ROI, to quantify Ca^2+^ activity in the FOV more accurately.

#### Microglial morphological analysis

The time-lapse stack was aligned offline using the Template Matching plugin of ImageJ Fiji to correct for xy motion artifacts. The image stack of 30 seconds in different brain states was compressed into one frame to analyze the morphological characteristics of microglia. Subsequently, the microglia morphology was reconstructed and quantified using Imaris 9.5 software. The definitions of microglial process length, area, and branch points remain consistent with our previously described^14^.

#### Adenosine signal analysis

Raw data were binned into 1 Hz, followed by background autofluorescence subtraction. ΔF/F_0_ were subsequently calculated, utilizing the fluorescence signal itself. F_0_ was defined as the average value of the lower 25th quartile fluorescence signal within each 60-second window of the recorded fluorescence signal. To enhance comparability across different brain states, wake and NREM states with a duration exceeding 30 seconds were selectively chosen for quantifying ADO signal changes. Mean ΔF/F_0_ of

ADO signals in different states within one session are used for comparison. Microglial Ca²⁺ and ADO signals were represented as z-score in representative long-term recordings (Fig. 1b; Fig. 2b). To quantify the ADO signal during brain-state transitions, the ΔF/F_0_ was further normalized^41^ (Fig. 2d).

#### EEG/EMG data analysis

All signals were digitally sampled at a rate of 128 Hz. EEG and EMG signals are processed using bandpass filtering (EEG, 0.1–35 Hz ; EMG, 10–100 Hz). Raw EMG signal passed 50 Hz digital notch filter. Signals were converted by using fast Fourier transform (FFT; FFT point, 256). In addition, typical oscillations in EEG were defined, such as delta (0.5–4 Hz), theta (4–8 Hz), spindles (8–12 Hz), and spindle (12–15 Hz). SleepSign software (Kissei Comtec) was used to score wake, NREM, and REM sleep states in 5-second epochs according to standard criteria^42^. At the same time, combined with behavioral videos, artificial correction was carried out for the states with obvious misjudgments in individual epochs.

#### Behavioral analysis

Behavioral videos were employed to differentiate behaviors such as locomotion, quiet wakefulness, and grooming. Locomotion was characterized as the horizontal movement process. Quiet wakefulness was conceptualized as an aroused state in which the mouse remained quiescent but awake. Sleep was ascertained through behavioral observations and electromyography recordings.

### Statistical analyses

All statistical analyses were conducted using GraphPad Prism 8 (GraphPad Software). For all tests, a two-tailed approach was employed to assess significance. Significance levels were denoted as follow: \**P* < 0.05, \*\**P* < 0.01, and \*\*\**P* < 0.001. Normality tests were performed on all data sets prior to further analysis. To determine the significance of differences between groups, appropriate parametric or non-parametric tests were utilized based on the data distribution and experimental design. Statistical details are provided in the figures and figure legends. All data are presented as the mean ± s.e.m.

## Supporting information

manuscript

## Acknowledgements

We thank Tong Zhu (Raygenitm Biotech Co., Ltd), Ying Guo, Yizhe Wei, Zhouyuan Liu and Xu Xie (Transcend Vivoscope Bio-Technology Co., Ltd) for comments on the optics and biological experiments; Dong Zhang, Jiaying Han and Fangxu Zhou (Peking University) for data processing; and the Nanjing Brain Observatory for data processing services. The work is supported by grants from the National Science and Technology Innovation 2030 Major Program (2021ZD0202205, 2022ZD0211903, 2021ZD0204002, and 2021ZD0204000), the National Natural Science Foundation of China (32293210, 92157105, 31971158, 81827809, 81827805, 82130060, and 61821002), CAMS Innovation Fund for Medical Sciences (2019-I2M-5-054), National Key Research and Development Program (2018YFA0704100 and 2018YFA0704104), Jiangsu Provincial Medical Innovation Center (CXZX202219), Collaborative Innovation Center of Radiation Medicine of Jiangsu Higher Education Institutions, Nanjing Life Health Science and Technology Project (202205045), and Key Core Technology Research Project for Nanjing Enterprise Academician Workstation.

## Author contributions

X.G. and H.C. conceived and supervised the project. Z.Z. and X.G. designed and supervised the biological experiments. Z.Z., X.G. and X.C. performed the experiments and led data analysis and figure-making; Z.Z. wrote the manuscript under the supervision of H.C., X.G., J.Y. and L.Z. All authors participated in the discussion, data interpretation and conception.

