## Supplementary material for "Microglia stabilize sleep homeostasis via adenosine A_3_ receptor signaling": manuscript

**Supplementary information**

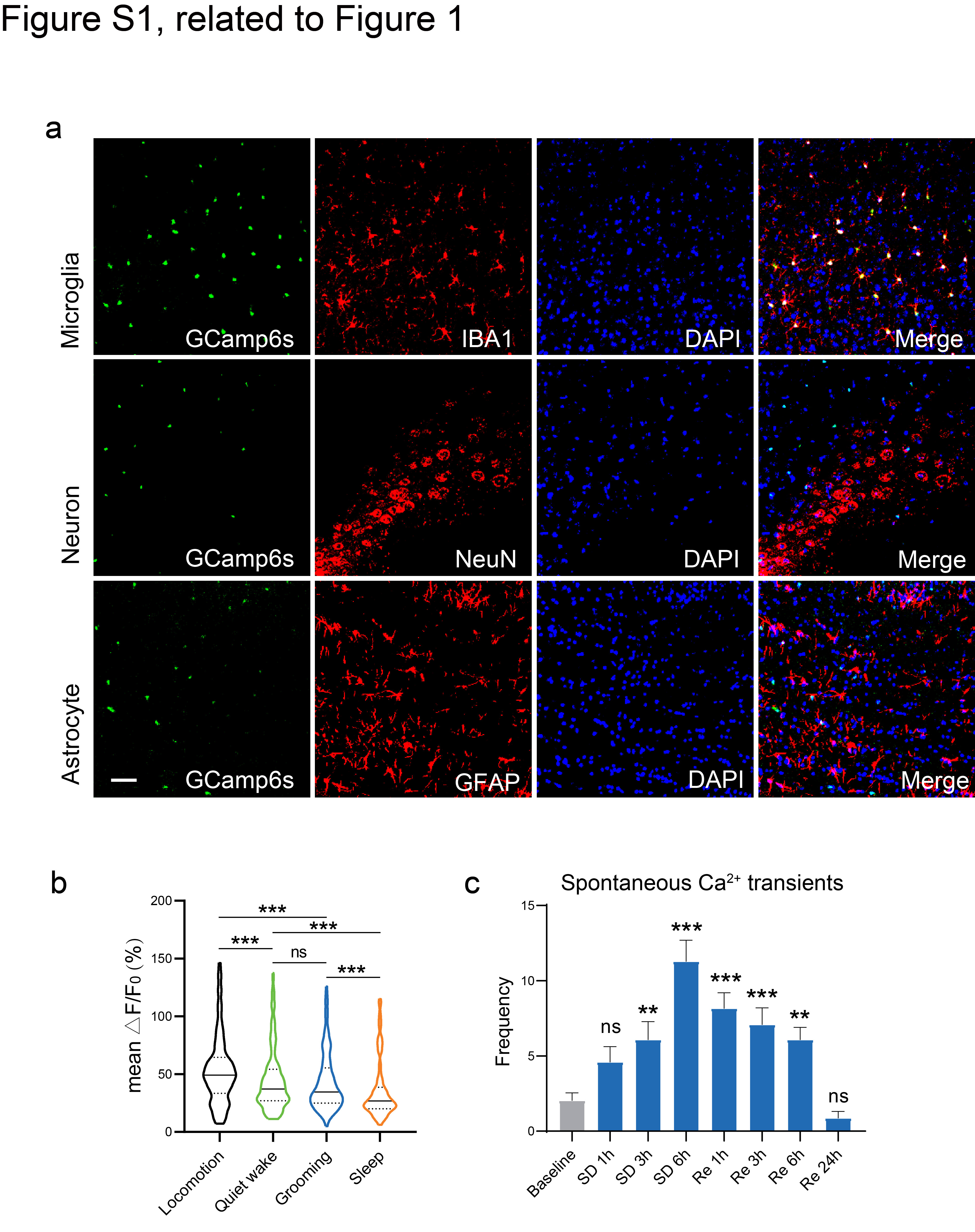

**Figure S1. Long-term microglial Ca^2+^ imaging in freely behaving mice. Related to Figure 1.**

**a.** Immunohistochemical verification of GCaMP6s expression in microglia (IBA1), but not neurons (NeuN) or astrocytes (GFAP). Nuclei were stained with DAPI. Scale bar, 200 μm. **b.** Mean Ca^2+^ activity of microglia under different behavioral conditions. Friedman test with Dunn’s post-hoc test was used. n = 166 cells from 6 sessions of 4 mice. **c.** The number of spontaneous Ca²⁺ transients in microglia recorded over a 15-minute period during sleep deprivation (SD) and recovery sleep (Re). n = 40 cells from 3 mice. unpaired t test. ***P*<0.01, ****P*<0.001. ns, not significant.

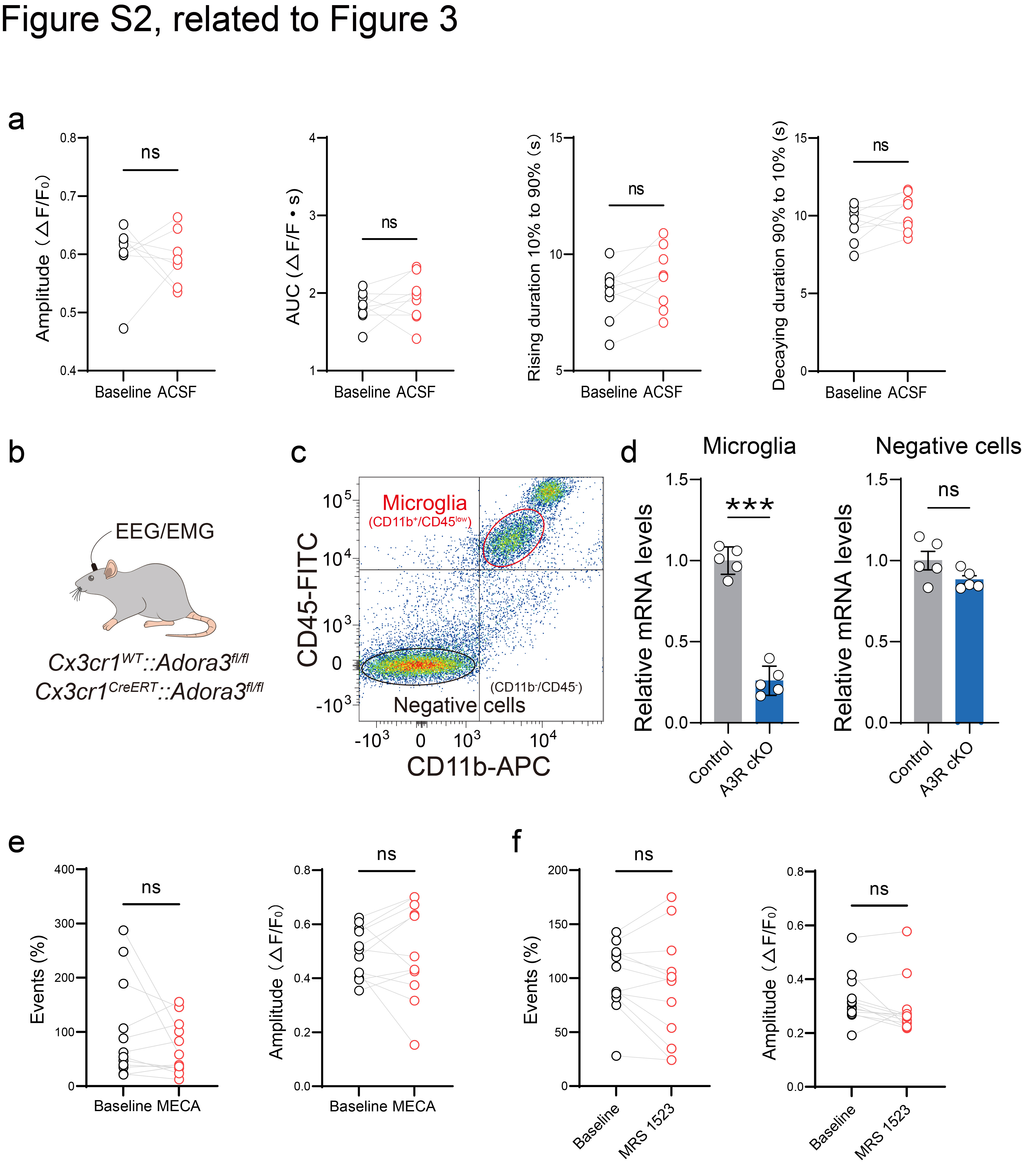

**Figure S2. A_3_R regulates microglial Ca^2+^ activity in microglia-specific A_3_R conditional knockout mice. Related to Figure 3.**

**a.** The amplitude, area under the curve (AUC), and duration of Ca^2+^ activity during microglia processes in the field of view (FOV) were analyzed in 5-minute recordings 30 min after ACSF administration. Data from the same FOV are connected by lines. n= 9 cells from 3 mice. Wilcoxon test (event, amplitude, AUC, rising duration, and decaying duration), two-tailed paired t test (duration). **b.** Experimental paradigm: cortical microglia were isolated one month after tamoxifen induction. *Cx3cr1^WT^::Adora3^fl/fl^* mice served as littermate controls. **c.** CD11b^+^/CD45^low^ cells (microglia) and CD11b^-^/CD45^-^ (negative cells) were collected by FACS. **d.** Quantitative RT-PCR showing reduced A_3_R mRNA expression in microglia from A_3_R cKO mice relative to control mice. Unpaired t test was used. n = 5 mice per group. **e-f.** The number of events and amplitude of Ca^2+^ activity during microglia processes in the FOV were analyzed in 5-minute recordings 30 min after Cl-IB-MECA (d) or MRS1523 (e) administration in microglial A_3_R cKO mice. Data from the same FOV are connected by lines. ****P*<0.001. ns, not significant.
